# Skull base invasive low-grade meningiomas, a distinct genetic subgroup: A microarray gene expression profile analysis

**DOI:** 10.1101/371716

**Authors:** Jun Sakuma, Masazumi Fujii, Yugo Kishida, Kenichiro Iwami, Keiko Oda, Kensho Iwatate, Masahiro Ichikawa, Mudathir Bakhit, Taku Sato, Satoshi Waguri, Shinya Watanabe, Kiyoshi Saito

## Abstract

**Introduction:** Meningioma is the most common adult primary brain tumor originating from meningeal coverings of the brain and spinal cord. Commonly, World Health Organization (WHO) grade-I meningiomas are slowly growing and surgically curative, some present with clinically aggressive behavior, invading the skull base bone and soft tissues by extending into the extracranial spaces.

**Methods:** To detect the genetic background of the Skull Base Invasive Low-grade Meningioma (SBILM), we conducted a comprehensive analysis of gene expression was conducted on 32 meningioma samples.

**Results:** The cluster analysis of the gene expression profile demonstrated a distinctive clustering pattern of the SBILM. Based on the clinical behavior and the microarray findings, they might be a distinct subgroup of meningiomas.

**Conclusion:** Further studies on characterization of genes specifically expressed by the SBILM could lead to the development of diagnostic tools, differentiating it from other WHO grade-I meningiomas and assist in the appropriate management and follow-up strategy, and open the door for development of pharmacological therapies.

## Introduction

Meningioma is defined as a group of mostly benign, slow-growing neoplasms that most likely derive from the meningothelial cells of the arachnoid layer. It is the most common tumor of the central nervous system (CNS), which account for approximately 36.4 % of all primary intracranial tumors and 53.4% of all non-malignant primary brain tumors in the USA[26]. Of meningioma with documented WHO grade, 81.3% of meningioma were WHO grade-I, 16.9% were WHO grade-II, and 1.7% were WHO grade-III[31].

Although most meningiomas grow slowly and toward the intradural space, some tumors exhibit higher-grade behavior in the form of recurrence and adjacent structures invasion[6]. Some WHO grade-I meningiomas do invade the skull base bone and even extend to the extracranial spaces. Those extending into the infratemporal fossa have an aggressive nature and invade bony floor of the middle fossa, skeletal muscles, nerves, and mucous membrane[34].

The term “invasive meningioma” has usually been linked to high- grade meningiomas, aiming at the brain parenchymal infiltration by the tumor cells[14,29]. Regarding skull base invasive meningiomas, the significant scientific work was conducted to study the surgical and radiotherapeutic strategies[15,19,37,42]. A little effort was made to study the biological background of Skull Base Invasive Low-grade Meningioma (SBILM).

Medical sciences had witnessed a significant improvement in the era of “genomic medicine”[10]. The combination of better biomarkers, more defined phenotyping and genomic/transcriptomic/proteomic analyses eventually allowed real progress in the diagnosis and management of currently neurological diseases. For the last 20 years, advanced technology has allowed us to measure levels of gene expression for many or all genes of an organism, and this has revolutionized our ability to screen the effect of genetic and environmental perturbations[27]. Meningioma was not an exception; in the last two-decade several papers were published regarding methods and reviews of the molecular biological profile of meningiomas[2,4,5,22,23,33,38,43,47,49]. The efforts of some of these studies helped in identifying unique molecular expression correlated with the WHO grades.

In our study, we hypothesized that SBILM possesses a distinct genetic expression profile than the rest of WHO grade-I tumors. Thus, we tested the genetic expression of tumor samples using microarray transcriptomic analysis system.

## Methods

### Sample collection and anonymization

Our definition of SBILM is based on the preoperative magnetic resonance images (MRI) and/or intraoperative findings. We included any WHO grade-I meningioma that invades the skull base bone with or without extension to the extracranial spaces and infiltration of other tissues such as muscles, nerves, and other extracranial structures.

Before surgery, written informed consents were obtained from each of the participants or their legal representative after providing them with information about the design and contents of the study both verbally and in written form. The study was reviewed and approved by the local ethics committee (No. 1953).

A cohort of random 32 patients with different forms of meningioma was included. Preoperative location of the tumor was documented based on MRI scans. Tumors locations were variable including meningioma of skull base and other sites. Tumor samples were surgically resected at our Institute between April 2009 and May 2010. Patients were 13 men and 19 women, and age ranged from 19 to 85 years (mean 61.2±14.8). Patient and tumors sample characterization are explained in Table 1.

At surgery, a tumor sample was divided into two parts: one was immediately frozen in liquid nitrogen for microarray study purpose, and the other was fixed for histopathological analysis. Tumor samples were anonymized by an assigned independent research personnel, who kept the connection table with the outboard recorder.

Diagnosis and classification of histopathological subtypes were based on WHO standard diagnostic criteria. Among 32 meningiomas, 25 cases were diagnosed as WHO grade-I, six as grade-II and one as grade-III (Table 1).

### Microarray transcriptomic analysis

#### Preparation of Poly A (+) RNA from clinical samples

Lysates of clinical tumor samples were subjected to total RNA extraction, followed by poly(A)+ RNA isolation with a MicroPoly(A)Purist™ Kit (Ambion Inc., TX, USA). The poly(A)+ RNA was divided into aliquots of 2 micrograms with precipitation with ethanol and stored at −20°C.

#### Microarray analysis

Synthetic polynucleotides (80-mer) representing 30913 species of human transcripts (MicroDiagnostic, Tokyo, Japan) were arrayed by using a custom arrayer. Two micrograms of poly(A)+ RNA was labeled with Cyanine 5-dUTP or Cyanine 3-dUTP (PerkinElmer Inc., MA, USA). Human common reference RNA was prepared by mixing equal amounts of poly(A)+ RNA extracted from 22 human cancer cell lines to reduce cell type-specific bias of expression. Labeling, hybridization, and wash of microarrays were performed using a Labeling & Hybridization Kit (MicroDiagnostic, Tokyo, Japan) according to the instructions of the manufacturer. Signals were measured with GenePix ^®^ 4000A scanner (Axon Instruments Inc., CA, USA) and then processed into primary expression ratios (ratio of the Cyanine-5 intensity of each cell line to the Cyanine-3 intensity of the human common reference RNA). Each ratio was normalized by multiplication with the normalization factor using the GenePix^®^ Pro 3.0 software (Axon Instruments Inc., CA, USA). The primary expression ratios were converted into binary log (log_2_) values (designated as log ratios).

#### Microarray data analysis

Data processing and hierarchical clustering analysis were performed using an MDI gene expression analysis software package (MicroDiagnostic, Tokyo, Japan). To identify genes demonstrating significant changes in expression, the t-test was performed between the invasive and non-invasive groups (p<0.001). Among the extracted genes, we further selected those genes that exhibited differences greater than 1.0 between the mean averages of binary log ratios for the two sample groups.

## Results

Of all 32 samples, six had skull base invasion. Five out of the six samples were WHO grade-I, thus were labeled as SBILM (Table 2). To focus on the altered gene expression between the SBILM samples and other samples of the WHO grade-I meningiomas, we did a hierarchical clustering analysis was conducted first to the 25 samples of WHO grade-I. First, genes with binary log values = 0 were removed. Then, statistically, significant genes were extracted using t-test (p < 0.001). Further, we selected genes with binary log value more than the value of 1 which represent a more robust alteration of expression. The result was a data set of 25 samples (WHO I) where SBILM showed a distinctive cluster of 36 genes (Fig.1).

**Fig.1.**
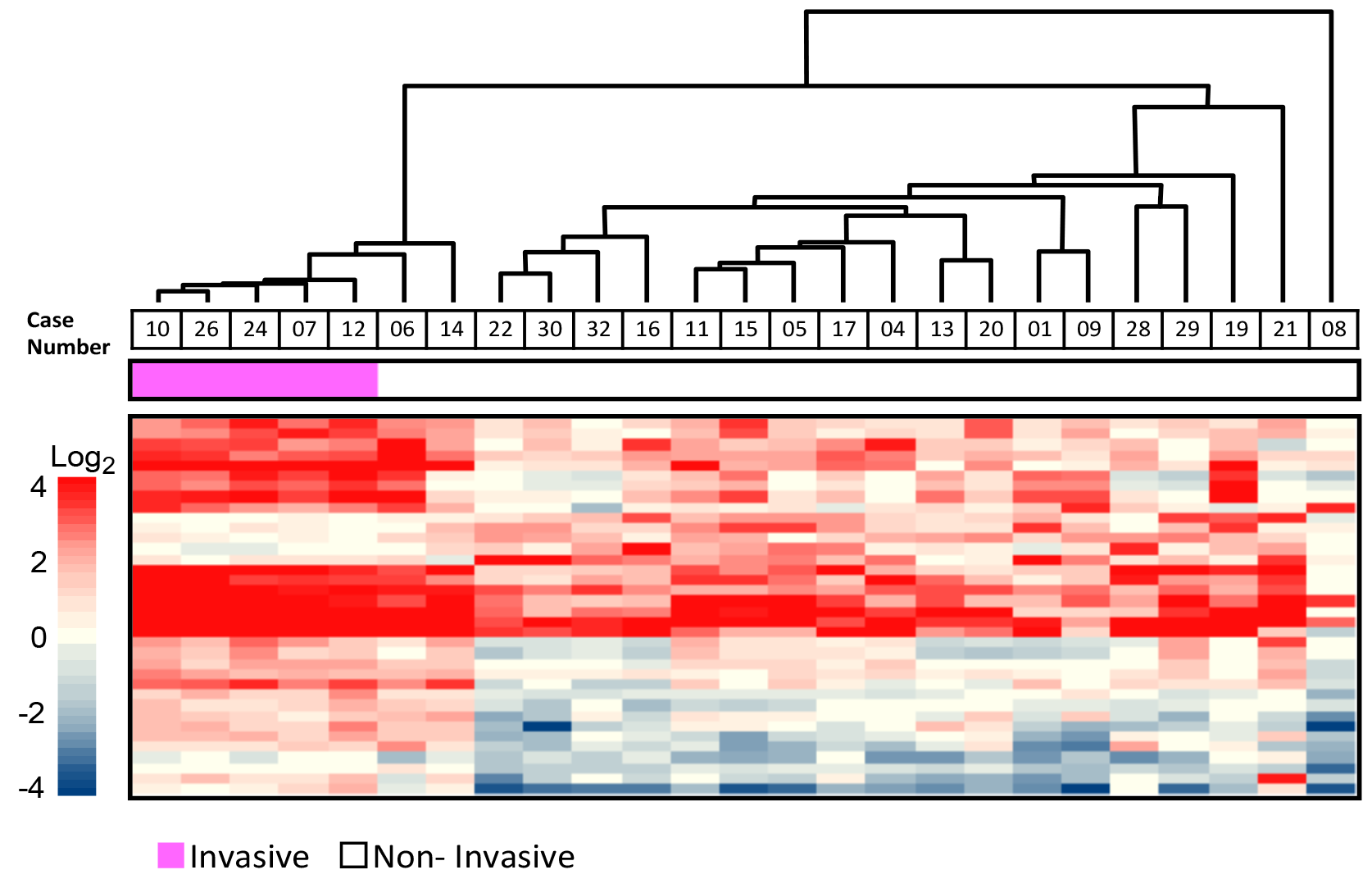
Dendrogram showing results of cluster analyses and heat map representation of the 25 WHO grade-I samples. SBILM are represented by magenta color in the second bar. Genes with < 1 log_2_ values were omitted. Statistically significant genes were extracted using t-test (p<0.001)

To confirm that this gene cluster pattern is characteristic to the SBILM we did hierarchical clustering analysis including all 32 samples with the same analytic steps. The SBILM showed the same distinctive gene cluster in comparison to the other samples (Fig. 2). Two other specimens (case No. 6 & 14) of WHO grade-I meningioma shared the same clustering pattern but at the time of diagnosis did not show skull base invasion (Fig. 1 & 2).

**Fig.2.**
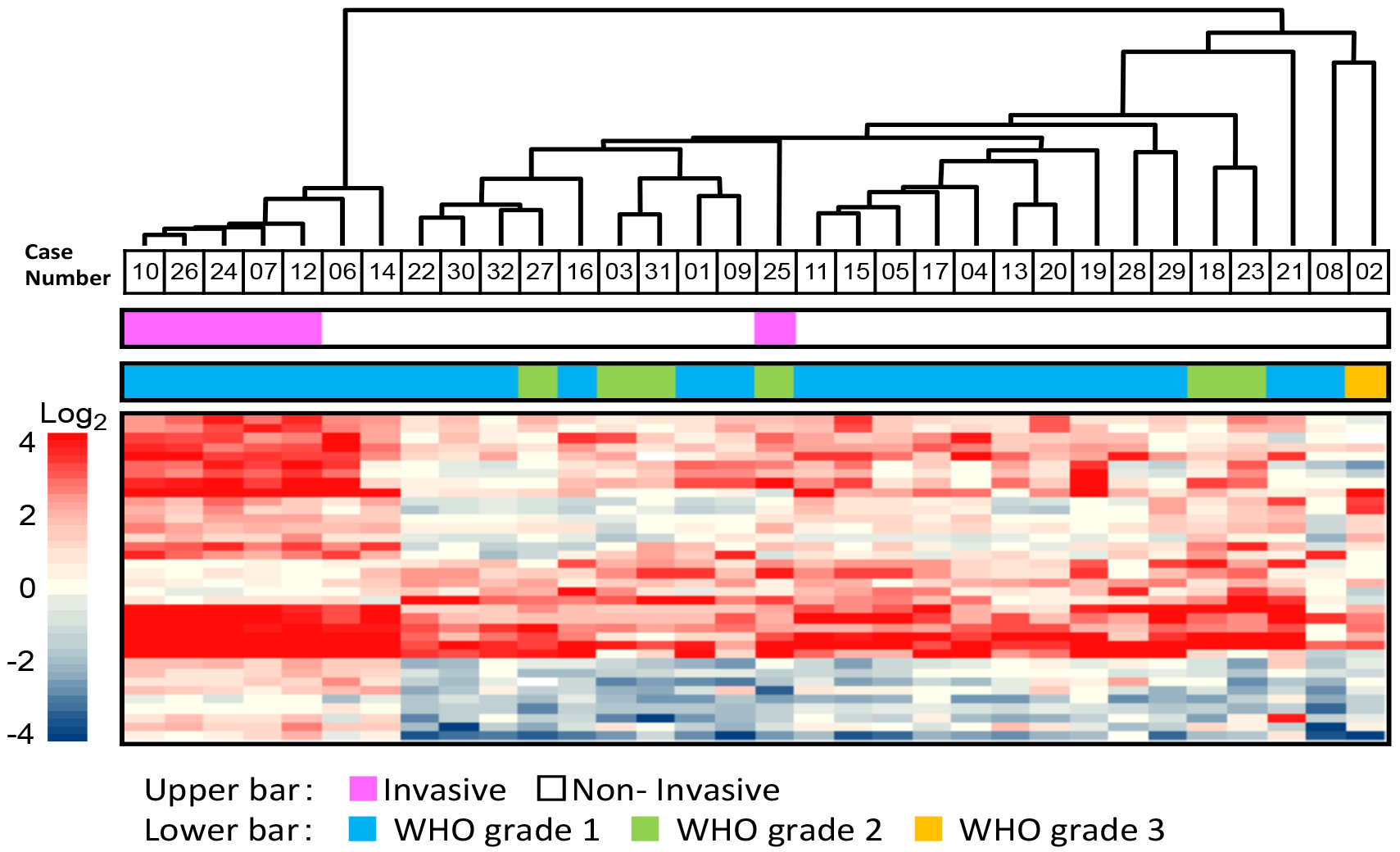
Dendrogram showing results of cluster analyses and heat map representation of the whole 32 samples. Skull base invasive meningiomas are represented by magenta color in the second bar. Genes with < 1 log_2_ values were omitted. Statistically significant genes were extracted using t-test (p<0.001). SBILM showed a different cluster pattern than the rest of the samples

All Five SBILM cases invaded skull base bone and extended into extracranial tissues (Illustrative cases Fig. 3 & 4) and shared a similar genetic expression. The extracranial invasion sites were the infratemporal fossa (2 cases), the paranasal sinuses (3 cases), clivus (1 case) and the orbit (2 cases).

**Fig.3.**
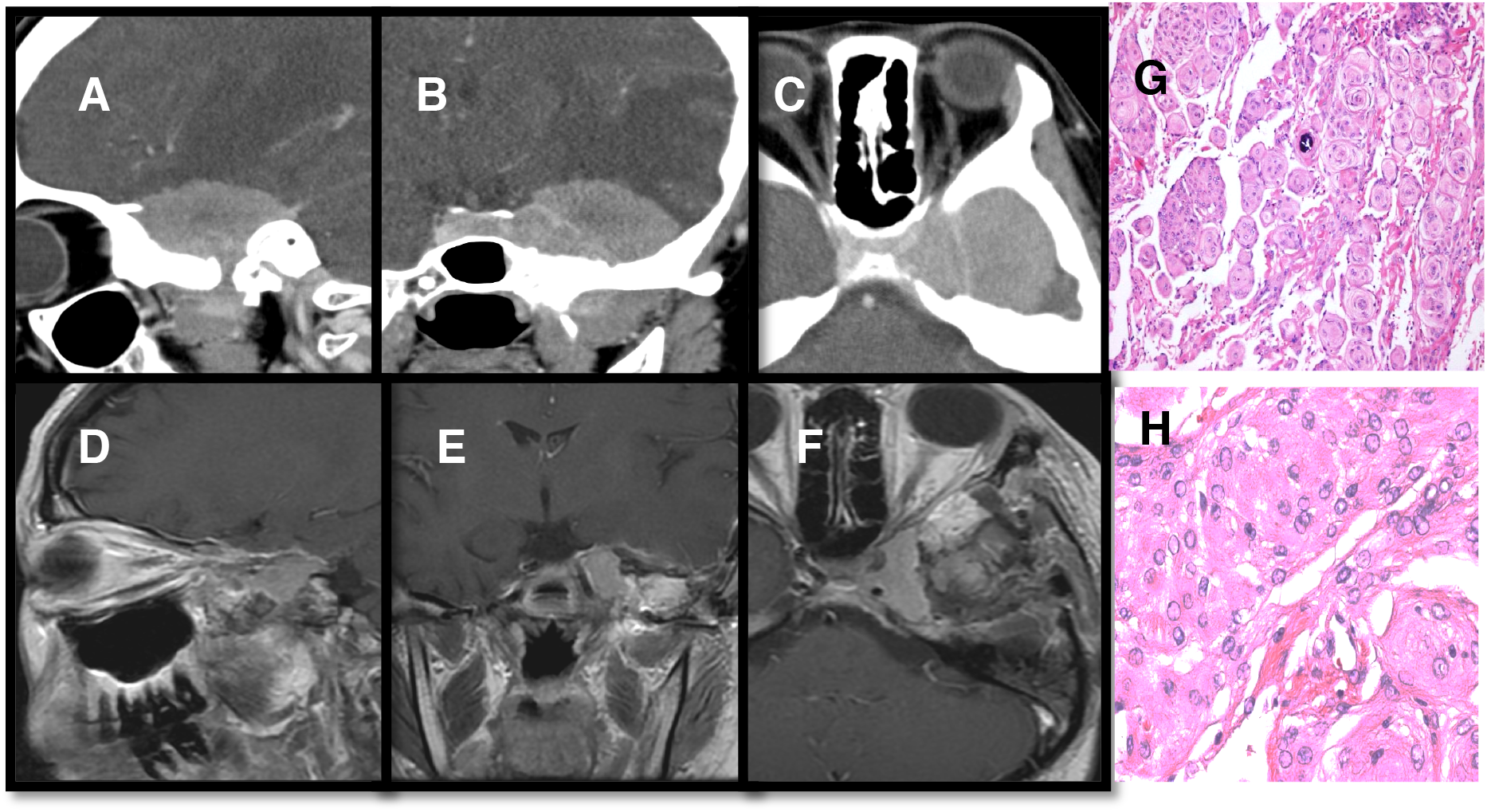
Case No. 24: A 29 years old female, with history of 4 months headache and facial numbness. CT brain Preoperative (A-C) showed left middle fossa meningioma with invasion to the cavernous sinus, skull base bone and soft tissue of the left infratemporal fossa. Intraoperative, the tumor in cavernous sinus was left. Otherwise the rest of the tumor was totally removed (D-F). Histopathology was WHO grade-I with MIB index < 1% but wasn’t possible to designate to a specific histological group (G, H)

**Fig.4.**
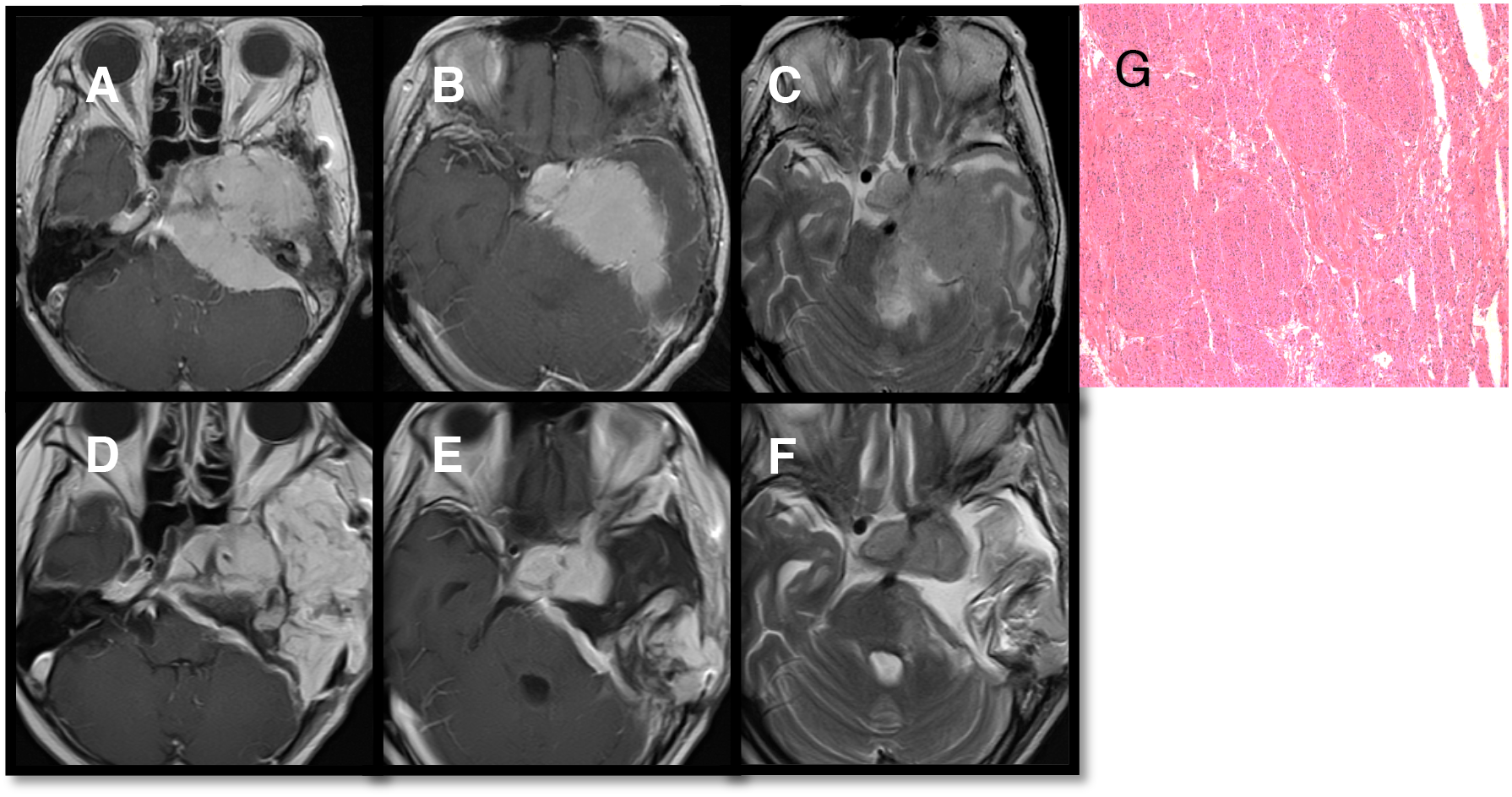
Case No. 12: A 67 years old female with recurrent skull base meningioma 13 years after the first subtotal resection. MRI demonstrated an invasive meningioma involving the cavernous sinus, the pituitary gland, the petrous bone, and the infratemporal fossa (A-C). Using a zygomatic combined petrosal (trans-labyrinthine) approach; the tumor was partially removed (D-F). Histopathological diagnosis was a meningothelial meningioma, WHO grade-I with MIB index < 1% (G). Remnant tumors in the cavernous sinus and pituitary fossa, and a tumor adhered to the brainstem was treated with gamma-knife radiosurgery

## Discussion

The WHO classification of meningiomas is widely used and is still primarily based on morphological parameters. More than 90% of all meningiomas are slow-growing tumors of WHO grade-I, and total excision of the tumor could achieve cure in most cases[26]. However, the use of the WHO classification as a tool to assess the clinical course and outcome has faced criticism. It became clear that the histopathological diagnosis alone can’t provide enough knowledge about the disease course and behavior. Efforts to improve the methods of tumor’s diagnosis and classifications were introduced especially after the significant achievements in tumors’ molecular biology field[9,18].

Despite their benign histopathological appearances, some of the “benign meningiomas” behave aggressively, invading the dura, dural sinuses, skull, and extracranial soft tissues and even the skin[3,32]. Notably, in the middle cranial base and sphenoid wing zones, an invasive growth to the extracranial spaces is more frequently seen. Although skull base meningiomas with bony invasion and extension to the extracranial spaces possess aggressive nature and harder to be completely resected, bone invasion or extracranial extension have not been considered as a diagnostic criterion of higher grade meningiomas.

Attempts have been made to early predict invasive meningiomas. There are several reports had investigated the invasiveness of meningioma and demonstrated its association with miRNA-18a, methylation of Werner syndrome protein, c-mET and hepatocyte growth factor, fatty acid metabolism-associated proteins or E-cadherin or β-catenin[21,24,25,44,50]. Schittenhelm et al. observed an increased secreted protein acidic and rich in cysteine expression is associated with the invasive phenotype in meningiomas, regardless of histological tumor grade[40]. Multiparameter analysis by Gay et al. using combined enzyme-linked immunosorbent assay identified Thrombospondin 1 (THBS1), as an overexpressed protein in infiltrative/invasive meningiomas. THBS1 is a protein that belongs to the group of extracellular matrix proteins and has been reported to control adhesion and de-adhesion between cells[14]. However, those studies investigated invasion of brain parenchyma by high-grade meningiomas, not the skull base invasion by low-grade ones. Beside parenchymal invasion, another group of studies focused on meningioma’s recurrence. The MIB-1 and proliferating cell nuclear antigen were found to predict recurrence, however, they did not predict invasive feature[45]. Another study showed that S100A5 protein could play a role in the recurrence of totally resected WHO grade I meningiomas[17]. Perez-Magan et al. in their study defined a differential gene expression pattern that distinguishes between primary and recurrent meningiomas. Most of these candidate genes are located at chromosomal regions whose loss has previously been associated with a higher risk of recurrence or malignant progression of meningiomas (1p, 6q, and 14q) [36]. Same author proposed candidate markers that could provide new information regarding the potential of these tumors to progress and/or recur and consequently may help predict clinical aggressiveness and improve patient management[35]. Ongaratti et al. reported that tumor grade was not significantly associated with recurrence or regrowth, which suggested another pathophysiological mechanism[30]. He identified a significant association between strong c-MYC expression and grades II and III meningiomas, underscoring the importance of the relationship between c-MYC expression and aggressive tumors. The latter report was not the only which tried to discover a link between the tumor’s grade, histopathology, and progression to the molecular biology of these tumors. Fèvre-Montange et al. in their results confirmed that several genes are differentially expressed in fibroblastic and meningothelial meningiomas[12]. Abedalthagafi et al. demonstrated that angiomatous meningiomas have distinct genomic features than any other WHO grade I meningioma[2]. Another study showed the expression of TERT mRNA was low in tumoral areas of low grade versus a sharply increased expression at the high-grade areas[1]. This finding apparently supports the presence of intratumoral heterogeneity. A recent whole transcript expression profiling study showed that Deleted in Colorectal Cancer might be a valid biomarker to identify those benign meningiomas at risk for progression[28]. The later study represents an example of how gene profile studies can assist in developing diagnostic markers for tumor progression.

Regarding skull base invasion, few reports were published. Wibom et al. examined the protein spectra in a series of meningiomas in search for a protein expression patterns that may distinguish between bone invasive and noninvasive meningiomas[48]. They found differences in protein expression between invasive and noninvasive, fibrous and meningothelial, meningiomas of WHO grade-I. Salehi et al. investigated immunoexpression of proteins implicated in meningioma skull base invasion[39]. They found higher expression levels of osteopontin and integrin beta-1 in invasive bone transbasal compared to noninvasive meningiomas. They also suggested that the molecular regulators of bone tropism, osteolytic activity, and vascular remodeling of meningiomas are dependent on anatomical location, with transbasal anterior skull base meningiomas showing a distinct differential expression pattern compared to spheno-orbital meningiomas. The later study might be the only study to our knowledge that examined the proteomic background of skull base invasive meningioma. To our knowledge, our study is the first to analyze the genetic expression of SBILM.

### Therapeutic strategy of the invasive meningiomas

The range of excision of meningiomas is strongly correlated with a rate of recurrence. The invasive meningiomas in the skull base, therefore, require an aggressive resection. Our surgical strategy for these kinds of tumors is as follows. We first remove the extradural parts of the tumor with bone, muscles, damaged nerves and mucosa with preservation of the functioning cranial nerves. Then we perform a resection of the intradural tumor, with its attaching dura. Skull base defects are reconstructed using either local flaps or vascularized free flaps. Stereotactic radiosurgery treats remnant tumors in the cavernous sinus or involving functioning cranial nerves. Even when histopathology results are benign tumors, we carefully observe the meningiomas with invasive features after surgery by more frequent imaging study than the none-invasive type.

### SBILM: Proposal of a new entity

In this study, we performed cluster analysis of gene expression profile, focused on genes expressed differently between the invasive and non-invasive skull base meningiomas. Our data included a group of 36 genes represented a unique clustering pattern with the SBILM.

A multiparameter analysis reported that meningiomas with brain invasion and those infiltrate the adjacent skull bone exhibited no distinguishable molecular features[14]. If this is the case, the entity of skull base meningioma invasive meningiomas might become even larger once including the brain-invasive meningiomas. Interestingly, in our sample, none of the SBILM did infiltrate the brain parenchyma (Table 2) (Fig. 4 & 5). Regardless of the small sample size, these results may suggest a molecular biological difference between SBILM and brain infiltrating meningiomas.

**Fig.5.**
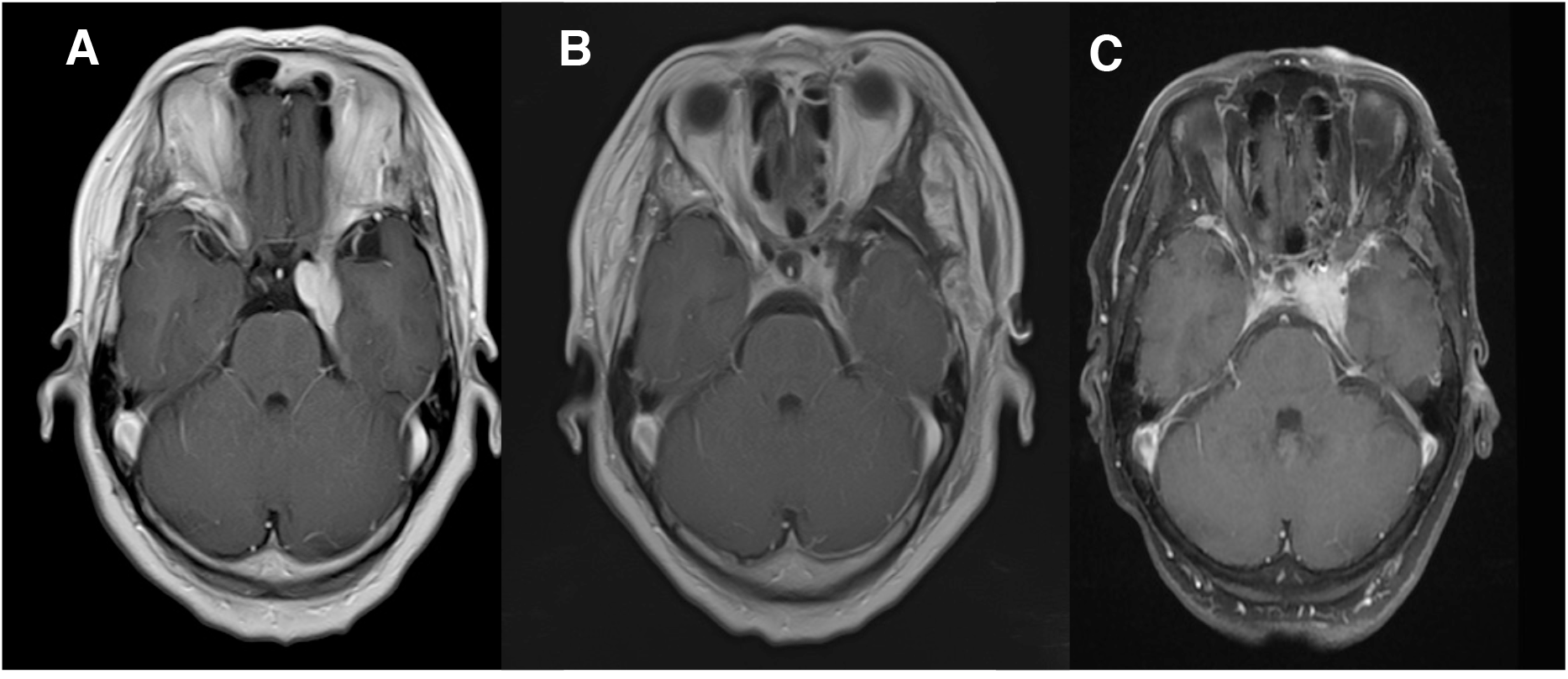
Case No. 14: A 65 years old female. (A) MRI brain was done and revealed a left cavernous sinus meningioma without apparent invasion into the skull base bone at the initial presentation. (B) Patient’s tumor was excised, and histopathology was a transitional meningioma, WHO grade-I. Gene expression microarray showed same gene cluster pattern of SBILM. (C) Patient had recurrence. She received stereotactic radiation

Genetic expression profile studies can provide tools for early detection of progressive tumors. If we predict invasive features of meningiomas in early stages even with mild or no clinical features, we can advantageously manage the patient after the first surgery. Furthermore, detection of over-regulated or down-regulated genes may assist in bringing personalized treatment for such tumors[13].

As mentioned in our results, two samples of non-invasive WHO grade I meningiomas (case No. 6 & 14) had the same genetic clustering pattern of the SBILM. We speculated that they “potentially” are SBILM and vigilant follow up might have been necessary. Unfortunately, we lost contact with one of the two cases after she moved to another region. The other patient (case No. 14) did follow up in another neurosurgery clinic for a period. She was 65 years old at her last visit. Her symptoms started with left ptosis. The patient was misdiagnosed as Tolosa- Hunt syndrome. The patient received medical therapy but clinically didn’t improve. MRI brain was done and revealed a left cavernous sinus meningioma (Fig. 6). The patient had surgery and tumor was excised. Histopathology showed a transitional meningioma, WHO grade-I. Gene expression microarray showed SBILM similar gene cluster pattern. The patient was offered careful follow-up in our clinic, but she requested to be followed in another clinic in her hometown. She had a recurrence of the tumor and received stereotactic radiosurgery. Unfortunately, follow-up was not feasible due to social reasons, and eventually, we lost contact with her.

Identification of tumors specific gene sequence might allow for the development of cancer-specific immune therapy or vaccine[16]. Vasudevan et al. did identify Forkhead Box M1 (FOXM1) as a master transcription factor for meningioma proliferation. They treated meningiomas cells with the FOXM1 antagonist FDI-6, had led to FOXM1 knockdown and pharmacologic inhibition of primary meningioma cell proliferation[46].

We have some limitations in this study. First, we have labeled the invasive meningiomas based on radiological and surgical findings. Some tumor might have already invaded the skull base at the micro levels, which cannot be detected by MRI or seen by naked eye. Careful follow-up of such cases is necessary since they might recur even after total resection.

Second, a tumor mass might possess a genetic heterogeneity[7,11,41]. Analysis of gene expression from a part of the tumor might not represent the entire molecular characteristics. It is necessary to perform analysis further on what extent heterogeneity of the relevant gene expressions exists and how it would have an impact on the clustering analysis, depending on the location of tissue samples.

Third, tumors may transform with genetic alterations when they grow or recur. Chou et al. in their study reported gene expressions in developing mouse brain exhibit a high level of spatial heterogeneity and temporal variation[8]. In the spatial dimension, they said the recurrent expression patterns that covered either globally (in most regions of the brain) global or locally (only in specific regions). If that were the case, the timing of tissue sampling would influence the results of the cluster analysis. Gene expression profiles of our cases should be compared when the tumors would recur in the future.

Fourth, technically speaking, the microarray is not perfect. Zhao et al. they reviewed the pros and cons of this technology[51]. According to their report, all genes will have an expression value in microarray due to background hybridization. Also, the microarray is less sensitive in the direct measurement of low abundant transcripts under different conditions. Furthermore, Microarrays are prone to "hybridization saturation" for highly abundant genes.

Last, in our study, we focused on the genetic expression and did not include the epigenetic analysis of the tumors. Epigenetics studies have proven to have a direct effect on the genetic expression process[20]. Altered DNA methylation, aberrant miRNA expression, and mutant epigenetic modifiers involved in histone and chromatin modifications being potential epigenetic markers of progression and recurrence of meningiomas[28]. Kishida et al. investigated the global methylation status of low-grade meningiomas with the aim of reclassification and compensation of the histological classification[22]. They reported the presence of subgroups with an aberrant accumulation of promoter methylation in WHO grade-I and II meningiomas. These alterations do correlate with high frequency of recurrence and may occur in the initial stage of histological changes such as malignant transformation. In another multicenter, retrospective analysis, Sahm et al. investigated genome-wide DNA methylation patterns of meningiomas[38]. They were able to identify patients at high risk of disease progression in tumors with WHO grade-I histology, and patients at lower risk of recurrence among WHO grade-II tumors

## Conclusion

The WHO grade-I meningiomas do include a group of cases which can demonstrate an aggressive behavior with aggressive skull base invasion. These SBILM tumors had shown a unique cluster analysis of gene expression in microarray transcriptomic analysis. This does represent a distinctive and unique entity of meningioma variants. SBILM technology allows us to detect those patients early to provide them with a cautious and vigilant follow up. Another benefit of genetic profiling is to allow for the development of target pharmacological therapies that can prevent tumor progression or recurrence.

## Funding

No funding was received for this research.

## Conflict of Interest

All authors certify that they have no affiliations with or involvement in any organization or entity with any financial interest (such as honoraria; educational grants; participation in speakers’ bureaus; membership, employment, consultancies, stock ownership, or other equity interest; and expert testimony or patent-licensing arrangements), or non-financial interest (such as personal or professional relationships, affiliations, knowledge or beliefs) in the subject matter or materials discussed in this manuscript.

## Ethical approval

The study was reviewed and approved by the local ethics committee (No. 1953).

## Informed consent

Before surgery, written informed consents were obtained from each of the participants or their legal representative after providing them with information about the design and contents of the study both verbally and in written form.

## Abbreviation

SBILM: Skull Base Invasive Low-grade Meningioma
MDI: Multiple Dataset Integration
UTP: Uridine-5’-triphosphate
miRNA: micro RNA
THBS1: Thrombospondin-1
TERT: Telomerase reverse transcriptase
FOXM1: Forkhead Box M1

